# NetTCR: sequence-based prediction of TCR binding to peptide-MHC complexes using convolutional neural networks

**DOI:** 10.1101/433706

**Authors:** Vanessa Isabell Jurtz, Leon Eyrich Jessen, Amalie Kai Bentzen, Martin Closter Jespersen, Swapnil Mahajan, Randi Vita, Kamilla Kjærgaard Jensen, Paolo Marcatili, Sine Reker Hadrup, Bjoern Peters, Morten Nielsen

## Abstract

Predicting epitopes recognized by cytotoxic T cells has been a long standing challenge within the field of immuno- and bioinformatics. While reliable predictions of peptide binding are available for most Major Histocompatibility Complex class I (MHCI) alleles, prediction models of T cell receptor (TCR) interactions with MHC class I-peptide complexes remain poor due to the limited amount of available training data. Recent next generation sequencing projects have however generated a considerable amount of data relating TCR sequences with their cognate HLA-peptide complex target. Here, we utilize such data to train a sequence-based predictor of the interaction between TCRs and peptides presented by the most common human MHCI allele, HLA-A*02:01. Our model is based on convolutional neural networks, which are especially designed to meet the challenges posed by the large length variations of TCRs. We show that such a sequence-based model allows for the identification of TCRs binding a given cognate peptide-MHC target out of a large pool of non-binding TCRs.

## Introduction

Cytotoxic T cells (CTLs) scan MHC class I-peptide complexes presented on the cell surface of nucleated cells. CTLs are able to recognize and kill infected or malfunctioning cells, e.g. cancer cells (1). Given the central role of the CTLs in the immune system, it is of paramount importance to understand the interaction between the T cell receptor (TCR) of the CTLs and their cognate peptide-MHCI targets. A peptide recognised in this context is referred to as a T-cell epitope.

The vast majority of all peptides that can be generated from a protein will not be presented by MHC molecules (2–4). Therefore, prediction of peptide-MHC binding is very useful to limit the number of peptide candidates when looking for potential T cell epitopes. Computational models have been trained successfully to predict peptide-MHCI binding, current state of the art methods include NetMHCpan (3, 4), NetMHCcons (5), NetMHC (6), the IEDB consensus method (7) and MHCflurry (8). Binding of peptides can be predicted with very high accuracy and precision for most human MHCI molecules (3).

However, not all MHC presented peptides are immunogenic. In order to predict which MHC restricted peptides do become T cell epitopes, the interaction between a TCR and its cognate target needs to be better understood. The TCR must be able to make contacts with the peptide as well as the MHC molecule to trigger an immune response. TCR and MHC interactions were reviewed by Gruta et al. (9). The focus of the here presented work is on the interactions between TCR and peptide.

Ample data are available linking peptides to the MHC molecules they bind, especially with the data obtained from mass spectrometry experiments (10–12). In contrast, there is much less data available linking specific TCRs to their cognate target. Recently developed high throughput sequencing methods are likely to change this situation and are already contributing increasing amounts of data (13, 14). Among those methods are the MIRA assay published by Klinger et al. (15) and the TCR barcoding technique published by Bentzen et al. (16). Additionally, two recent publications by Glanville et al.(17) and Dash et al. (18) have made more high throughput data available. Furthermore, these works describe clustering algorithms able to group TCRs by their epitope specificity. In particular, the work by Glanville et al. suggests that relatively simple sequence-based models can be used to classify and define specificity groups shared by TCRs and individuals. This is in line with earlier work by Roomp and Domingues (19). Several structure-based approaches for modelling the structure and interactions of the TCR:p:MHC system have likewise been proposed including structural modeling (20, 21) and structure based prediction of TCR:p:MHC interactions (22).

The IEDB (23) as well as the VDJdb (24) collect sequenced TCRs with known specificity published in peer reviewed articles, thereby providing a useful data resource to the community.

Here we seek, based on such data, to go beyond the work by Glanville et al. and present NetTCR, a method to predict the interaction between TCRs and peptides presented by HLA-A*02:01. NetTCR is based on convolutional neural networks (CNNs) and depends only on the amino acid sequences of the peptide and CDR3 region of the TCR beta chain as input. CNNs scan their input with convolutional filters that cover only short continuous parts of the input. These filters detect patterns and the network is then able to integrate the information from the patterns discovered by different filters throughout the input sequence. This type of model has been very useful in image classification (25), and recently also for handling sequence data of variable length for prediction of for instance protein secondary structure (26, 27), kinase phosphorylation (28) and subcellular location (29). CNNs are also ideally suited to deal with unaligned peptide and TCR sequences differing in length. The here presented model is available as web-server under http://www.cbs.dtu.dk/services/NetTCR/ and the underlying code can be downloaded here: https://github.com/mnielLab/netTCR.

## Methods

### Data set

A dataset of TCR beta chain CDR3 sequences and corresponding cognate peptide targets was downloaded from the IEDB (23) in April 2018. Only peptides presented by HLA-A*02:01 were selected. The training data consisted of 9015 unique data points, spanning 91 peptides and 8920 TCR sequences. Further, an additional dataset generated using the MIRA assay was kindly provided by Klinger et al. (15). This dataset consisted of 379 unique data points, spanning 16 peptides and 379 TCR sequences derived from 5 donors.

Since these data sets contain only positive interactions, negative data examples were generated by creating internal wrong combinations of TCRs and peptides, i.e. combining TCR sequences with peptides different from their cognate target. These combinations were made by extracting the list of peptide targets from the positive data set (keeping duplicates if a peptide was found to interact with multiple TCRs), and next pairing each TCR with a peptide different from the cognate target randomly drawn from this list of peptide targets. In this way, a data set with 50% positive and 50% negative data points was obtained.

To supplement the data sets with additional negative examples, eluted peptide ligands were retrieved from the IEDB, selecting only peptides derived from self (i.e human) proteins. Further, a set of 200,000 TCR CDR3 sequences from 20 healthy donors (30) was downloaded. Next, additional data was created by first replacing each TCR in the combined positive and negative mis-paired data sets three times with a random TCR drawn from the healthy donor TCR data set (proportions of additional negatives ranging from 2–5 were tested with limited variations in validation predictive performance, data not shown). Finally, the 3000 eluted ligands were paired with TCRs drawn randomly from the complete set of IEDB positives, IEDB mis-paired negatives, and additional negatives constructed from the TCRs of healthy donors. All the additional TCR peptide combinations were added as negatives to the data set, obtaining a final training data set consisting of 9012 (12%) positive and 66,102 (88%) negative TCR-peptide combinations.

To avoid model overfitting and overestimation of model performance, the entire data set was partitioned into 5 sets prior to model training. Prior to partitioning, TCR beta chain (TCRb) CDR3 sequences were compared to each other using blastp and TCRs sharing more than 90% sequence identity, determined by blastp, were kept in the same data partition. Otherwise data points were assigned to partitions randomly.

### Model

A CNN model was implemented to predict whether or not a given TCR is able to recognize a specific peptide. The input to the network was the amino acid sequence of the peptide and the CDR3 region of the beta chain of the TCR. Both sequences were encoded using the BLOSUM50 matrix as described earlier (31). Additionally N- and C-terminus start and stop signals were added to the TCR sequence and encoded using a vector containing only +/−0.1 for start and stop respectively. Peptide and TCR sequences were each processed by 100 convolutional filters of sizes 1, 3, 5, 7 and 9 amino acids (500 filters in total). The peptide and TCR convolutional layers were concatenated and processed by a second convolutional layer of 100 filters with size 1. Subsequently, global max pooling was performed to remove the sequence length dimension of peptide and TCRb CDR3 region. Global max pooling results were connected to a dense layer of 10 hidden neurons connected to the output neuron. Throughout the network, the sigmoid activation function was applied to all neurons. The network was trained using 5 fold cross validation with early stopping for 300 epochs. The weights were updated using the adam optimizer with a learning rate of 0.001. When additional negative examples were added to the training process, a subset of the negative examples was randomly selected in each training epoch to keep the frequency of positive data points at 50%. The validation error (to determine the early stopping epoch) however, was calculated on the full unbalanced data set.

All models were implemented in the Python programming language using the tensorflow library. The models were exclusively trained on the IEDB data, the MIRA data was used solely for performance evaluation purposes. Model performance was measured in AUC (area under the ROC curve, 0.5 corresponding to random predictions, 1.0 equals perfect predictions) or AUC10%, the partial AUC integrated upto a false positive rate of 10%.

### Model evaluation

#### Performance on a large set of TCRs

To investigate whether the model was able to identify the TCRs interacting with a given peptide out of a large set of TCRs, we identified 3 peptides (GILGFVFTL, GLCTLVAML and NLVPMVATV), all frequently occurring in both the IEDB and MIRA data sets. Each TCR in the MIRA data was paired with each of the three selected peptides and the pair was annotated depending on whether this interaction is positive (i.e. observed in the MIRA experiments) or negative. With this setup, the TCRs can be partitioned into two groups: 1) TCRs with a target among the three analyzed peptides and 2) TCRs without a target among the analyzed peptides. The data were predicted using the models described above (trained on the IEDB data, with and without additional negative data). Maximum prediction values of TCRs not targeting any of the 3 peptides were compared to the prediction values obtained for positive peptide-TCR pairs. Further, the predictions of a given TCR to all 3 peptides were ranked and the rank of the true binding peptide was extracted.

#### Performance on a set of randomly assigned TCRs

To provide a random baseline performance, a data set was generated by randomly assigning the TCR sequences of the original data to the peptides in the data set. Subsequently a model was trained on this random data set as described above.

#### V+J gene bias

To investigate how much of the models performance could be explained by a potential bias in V and J gene usage, models were trained on the center and N- and C-terminal regions of the TCRb CDR3 sequences. To train models on only the central part of the CDR3 sequence, the first and last two amino acids were replaced with X, corresponding to unknown amino acid. The influence of the N- and C-terminal regions of the CDR3 sequence was investigated using two approaches: 1) the TCRb CDR3 sequence represented only by the first and last two amino acids and 2) all amino acids of the TCRb CDR3 sequence except the first and last two are replaced by X, thus conserving information about the length of the original CDR3 sequence. Separate models were trained on these three representations of the TCR sequences and compared to a model trained on the complete CDR3 sequence.

#### Experimental validation

##### Ethical approval

All healthy donor material was collected under approval by the Scientific Ethics Committee of the Capital Region of Denmark, and written informed consent was obtained according to the Declaration of Helsinki.

##### Peptides and MHC monomer production

Peptides were purchased from Pepscan (Pepscan Presto) and dissolved to 10 mM in DMSO. UV-sensitive ligands were synthesized as previously described (32–34). Recombinant HLA-A*02:01 heavy chain and human β2 microglobulin light chain were produced in Escherichia coli. HLA heavy and light chain were refolded with UV-sensitive ligands and purified as described in (35). Specific peptide-MHC complexes were generated by UV-mediated peptide MHC exchange (33).

##### Generation of fluorescently labeled pMHC tetramers

MHC tetramers were assembled as described previously (36, 37) onto one of two fluorescently-labeled streptavidin (SA) conjugates: SA-phycoerythrin (PE) or SA-allophycocyanin (APC) (BioLegend, Nordic Biosite, Denmark). Tetramers binding one of the four peptides NLVPMVATV, GLCTLVAML, YVLDHLIVV and GILGFVFTL were labeled with PE while the remaining tetramers were labeled with APC (see Table S2 for full list of included peptides). Tetramers were stored at –20 °C in 5% glycerol (vol/vol) and 0.5% BSA (wt/vol).

Frequencies of antigen-specific T cells (Table S2) were determined using combinatorial encoding of pMHC tetramers (36, 37) or DNA barcode-labeled MHC multimers (16).

##### Peptide-MHC tetramer staining

Cryopreserved PBMCs from four healthy donors were thawed and washed in RPMI + 10% FCS. Cells were washed in a cytometry buffer (PBS + 2% FCS). 5×10^6^ cells were incubated, 15 min, 37 °C, with pooled PE and APC tetramers in a total volume of 100 μL (final concentration of each distinct pMHC, 23 nM). Next a 5× antibody mix composed of CD8-BV480 (BD 566121, clone RPA-T8) (final dilution 1/50), dump channel antibodies: CD4-FITC (BD 345768) (final dilution 1/80), CD14-FITC (BD 345784) (final dilution 1/32), CD19-FITC (BD 345776) (final dilution 1/16), CD40-FITC (Serotech MCA1590F) (final dilution 1/40), CD16-FITC (BD 335035) (final dilution 1/64) and a dead cell marker (LIVE/DEAD Fixable Near-IR; Invitrogen L10119) (final dilution 1/1000) was added and incubated 30 min, 4 °C. Cells were washed twice in cytometry buffer, resuspended in 100 μL cytometry buffer and sorted immediately.

##### Flow cytometry and cell sorting

Tetramer-stained cells were sorted on a FACSAriaFusion (Becton Dickinson) into tubes containing 100 μL PBS supplemented with BSA (0.5%), herring DNA (100 μg/mL) and EDTA (2 mM) (tubes were saturated with PBS + 2% BSA in advance). Using FACSDiva software, we gated on single, live CD8 positive and ‘dump’ (CD4, 14, 16, 19, and 40) negative lymphocytes and within this population sorted either all PE positive cells or all APC positive cells into separate tubes. The sorted cells were centrifuged 10 min, 5,000g, and the buffer was removed. The cell pellet was stored at −20 °C in a minimal amount of residual buffer (<20 μL). DNA was isolated using QIAamp DNA Micro Kit according to manufacturer’s instructions (Qiagen) and the TCRb chains were sequenced and processed at Adaptive Biotechnologies (Seattle, WA) using the ImmunoSEQ platform.

##### In silico interaction predictions

Binding to all four peptides was predicted for the PE (now referred to as positive) and the APC (negative) sorted populations (Table S3) using the TCRb CDR3 sequences and the model trained on a combination of IEDB and MIRA data sets with additional negative data. For a given TCR sequence the maximum scoring prediction was recorded and it was investigated whether a difference in these maximum prediction scores could be observed between the positive and negative TCRs.

## Results

We here present a machine learning model to predict TCR-peptide interactions based on only the peptide and TCRb CDR3 amino acid sequences. Training data was obtained from the IEDB, the model was evaluated on data generated with the MIRA assay (for details see materials and methods). Both datasets cover several peptides presented by HLA-A*02:01 but are dominated by the same 3 peptides (Figure 1A). Prior to training the model, the data were partitioned keeping similar TCRs in the same partition to limit redundancy between data partitions. Figure 1B shows all TCRs in the IEDB data color coded according to their peptide target (TCRs recognizing a peptide different from the above mentioned 3 abundant peptides are colored gray), lines connect TCRs sharing more than 90% sequence identity as determined by BlastP. This threshold resulted in clusters largely specific for one given peptide, indicating it is appropriate to reduce data redundancy.

A convolutional neural network architecture used to predict interaction between TCRs and their cognate target is shown in Figure 1C. The two convolutional layers combined with global max pooling enable training the model on peptides and TCR sequences of different lengths. Models were trained using 5 fold cross-validation as described in materials and methods.

**Figure 1:**
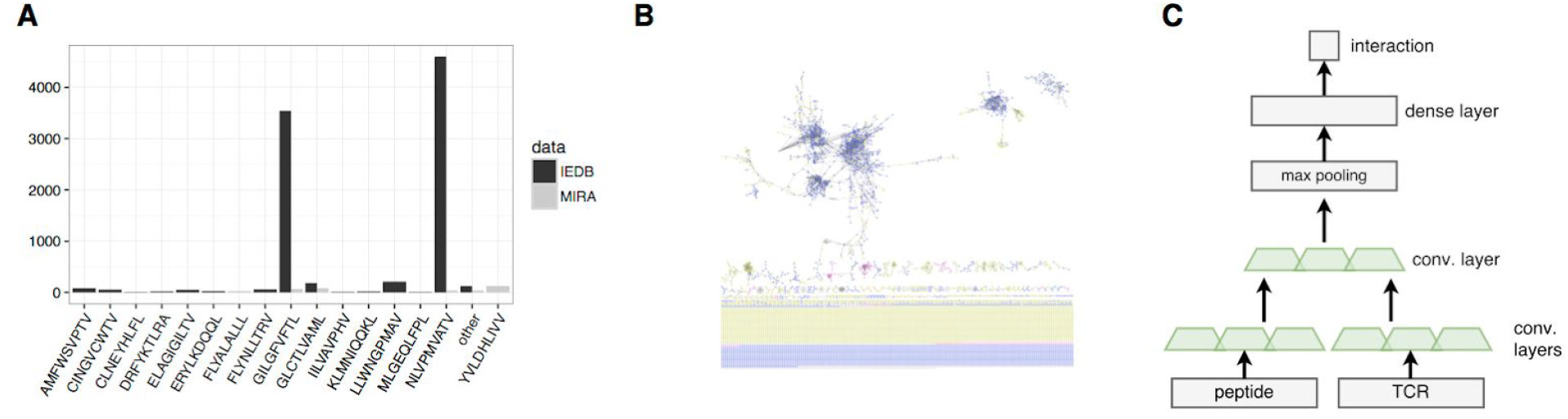
A) Frequency of peptides in IEDB and MIRA data sets. Only peptides that occur more than 10 times are shown, less frequent peptides are summarized in “other”. B) TCRb CDR3 clustering in the IEDB data set. The coloring corresponds to the TCRs peptide target, TCRs that share more than 90% BlastP sequence identity are connected C) Setup of the convolutional neural network.

### Model performance

One important application of our model would be to identify binding TCRs specific to one or more of the peptides from a large data set of irrelevant TCRs obtained, for instance, by repertoire sequencing. We simulated this task by selecting the three most common peptides in the IEDB (GILGFVFTL, GLCTLVAML and NLVPMVATV) which are also part of the MIRA data. Subsequently, we predicted binding of all TCRs in the MIRA data to each of those three peptides, using two different models: one trained on the IEDB data with internal negative data and another trained with additional negative data (derived from TCR sequencing projects and eluted peptide ligands, for details see methods). Subsequently we evaluated how the models could separate positive TCRs binding one of the three peptides from the negative TCRs.

Figures 2A-C give the results of this analysis, comparing the performance of the two models in terms of ROC, sensitivity and specificity curves. The AUC value of the model trained with additional negative data was slightly increased (0.697 to 0.727) compared to the model trained with only internal negative examples. As shown in Figure 2A, the AUC10% increased substantially to 0.48 from 0.27 with additional negative data. Figure 2B reveals that additional negative training data increased the specificity of the model while decreasing the sensitivity (Figure 2C).

**Figure 2:**
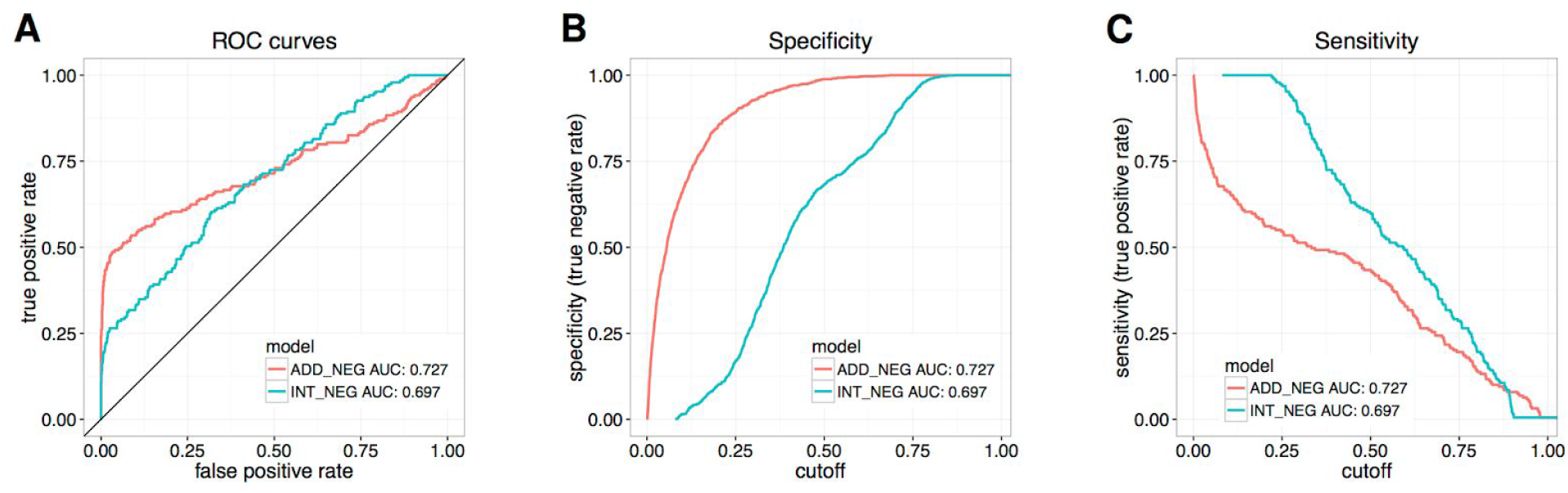
A) ROC curves, B) specificity and C) sensitivity for models trained on the IEDB data with internal negative data and additional negative data. Predictions were made on three common peptides shared between IEDB and MIRA data, combined with all MIRA TCRs.

Next, we investigated to what degree our model was able to identify the correct cognate peptide target for a given TCR. For each TCR binding to one of the three peptides included in the evaluation data set we predicted binding to all three peptides, and calculated the rank of the true cognate target. This rank is 1 if the cognate peptide target achieved the strongest predicted binding value among the three peptides. Figure 3A shows the histogram of these rank values for the two models. The model trained with additional negative data predicted the target peptide as rank 1 for 112 (59.3%) TCRs out of 189 compared to 90 (47.6%) for the model trained only on IEDB data. These results thus confirm the predictive power of the method in identifying the correct cognate target for a given TCR.

Apart from producing high predictions for the interaction between TCR and cognate target peptide, in order to be useful, a model is also required to make low predictions for negative TCRs with no cognate target among the peptides covered by the model. We tested to which degree this was the case for the models trained with or without additional negative data. For this we compared the highest prediction values obtained for TCRs without a target among the three selected peptides, to the predictions made for TCRs paired with their correct target. The result of this comparison is shown in Figure 3B and revealed that the model trained only on the IEDB data often assigned negative TCRs higher prediction values compared to observed peptide-TCR pairs. In contrast, for the model trained with additional negative examples, we found that the true cognate targets received higher median prediction values than the maximal predictions made for TCRs without cognate targets among the selected peptides. Similar observations were made when considering the predictions made for a specific peptide (Figure S1). Also in this case, the model with additional negative training data predicted noticeably lower values for negative compared to positive TCRs.

In conclusion, these results demonstrate that the model trained on additional negative data outperformed the model trained only on IEDB data in terms of assigning the highest rank to true target peptides. Further, this model was able to accomplish the task of finding few interacting TCRs out of a pool of many non-interacting TCRs, due to increased specificity.

**Figure 3:**
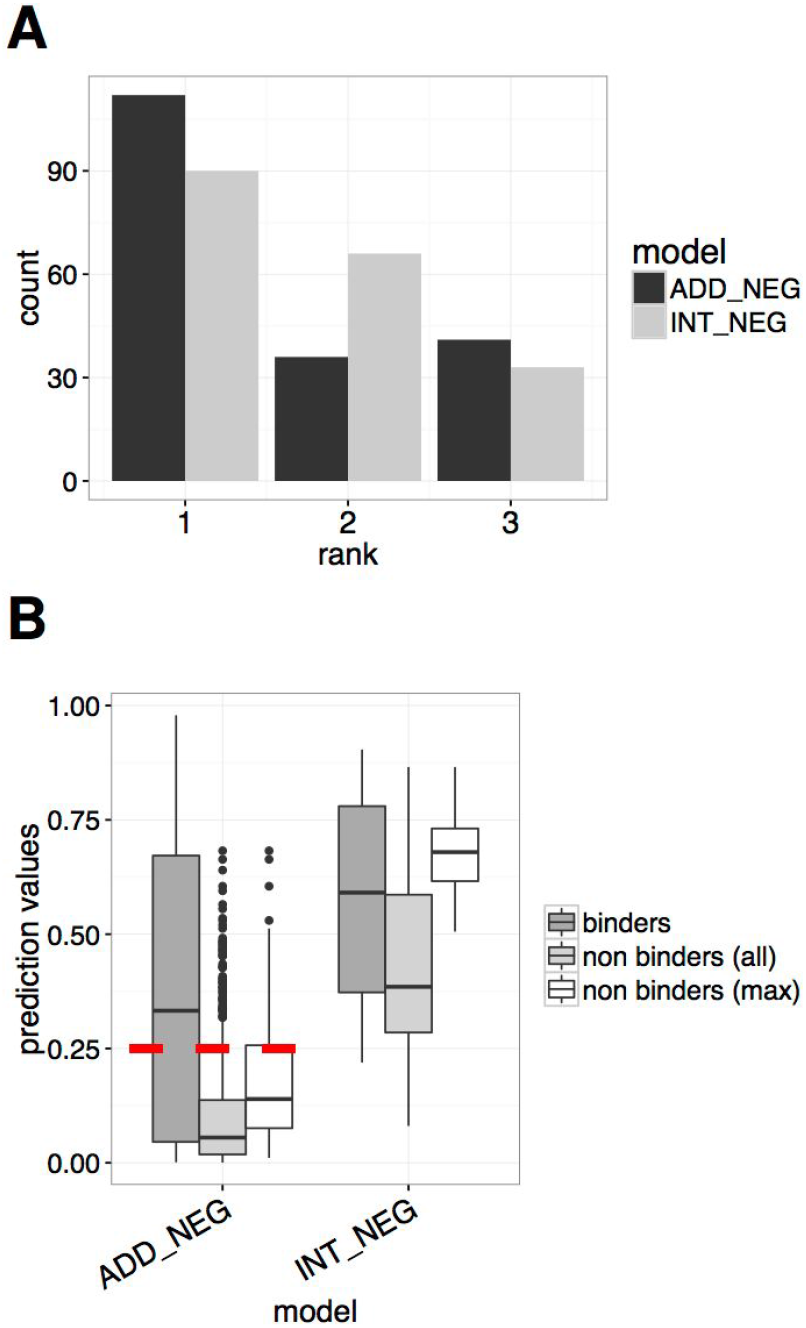
A) For the TCRs binding one of the 3 peptides in the evaluation set the rank of this peptide is shown (rank 1 = highest prediction, rank 3=lowest prediction). B) For TCRs binding to one of the three peptides in the evaluation set the prediction value to this peptide is shown. For TCRs not binding to one of the peptides in the evaluation set all prediction values are shown. Subsequently only the maximum prediction to an evaluation peptide is shown. The intention is to visualize if one can separate TCRs that have a cognate target among the peptides the model is trained on, from those that do not recognize any of the peptides in the model, based the model’s prediction values. The red line at 0.25 indicates such a classification threshold.

#### Performance on unknown peptides and randomly assigned TCRs

The IEDB data set is dominated by only 2 peptides, as shown above in Figure 1A. Still, we aimed to investigate if our model is able to learn general interaction rules that allow for accurate predictions for peptides not included in training. To get an estimate of the model’s performance on unknown peptides, we predicted the MIRA data and calculated the performance per peptide. The result of this analysis is shown in table S1 and demonstrated clearly that the performance on the peptides unique to the MIRA data was considerably lower than the performance on peptides shared with the training data. This indicates that the model has limited potential for extrapolating predictions to unknown peptides, most likely due to the limited amount of peptides in the training data.

To obtain a baseline performance and ensure that this would be random, we trained a model where all TCRs in the data set were randomly re-assigned to a peptide. As expected this model achieved a random performance of AUC=0.538 on the test set and AUC=0.4 on the external MIRA data.

#### Influence of V and J genes on model performance

The current data sets linking TCRs to peptide epitopes are small and further limited by being derived from a small number of donors. This could possibly lead to a bias in the V and J gene usage of the donors to which a model predicting peptide and TCR interaction might overfit. As the N- and C-terminal parts of the CDR3 sequence are defined mostly by the V and J genes, we tested the potential V and J gene bias in the data by comparing the performance of a model trained on the full TCRb CDR3 sequence to models trained only on the central part of the CDR3 sequence or the N- and C-terminal parts, (see figure 4A). For evaluating the amount of information captured in the terminal parts, we represented the TCRs as only the first two N and C terminal residues or with the central part of the sequence masked as X for unknown amino acid, thereby conserving information about the loop length of the original CDR3 sequence (for details see materials and methods).

Figure 4B shows the test set performance of the different models. Training only on the two N- and C-terminal amino acids of the CDR3 sequence resulted in a marked drop in test set performance (from AUC=0.676 to AUC=0.605), indicating there is not enough information contained in the terminal residues only to train a prediction model. When informing the model of the N and C terminal amino acids along with the loop length of the CDR3 region, a model can be trained with a test performance comparable to when using the complete CDR3 sequence (AUC=0.655). This is also the case when training only on the central part of the CDR3 sequence, masking the N- and C-terminal residue (AUC=0.664). However, when turning to the external MIRA evaluation set (see figure 4C), only models trained on the the complete or central part of the CDR3 sequence generalize well. This result suggests that models trained only on the N- and C-terminal regions of the CDR3 sequences likely overfits to the V and J gene distribution in the IEDB and therefore do not achieve good performance on the MIRA data set.

**Figure 4:**
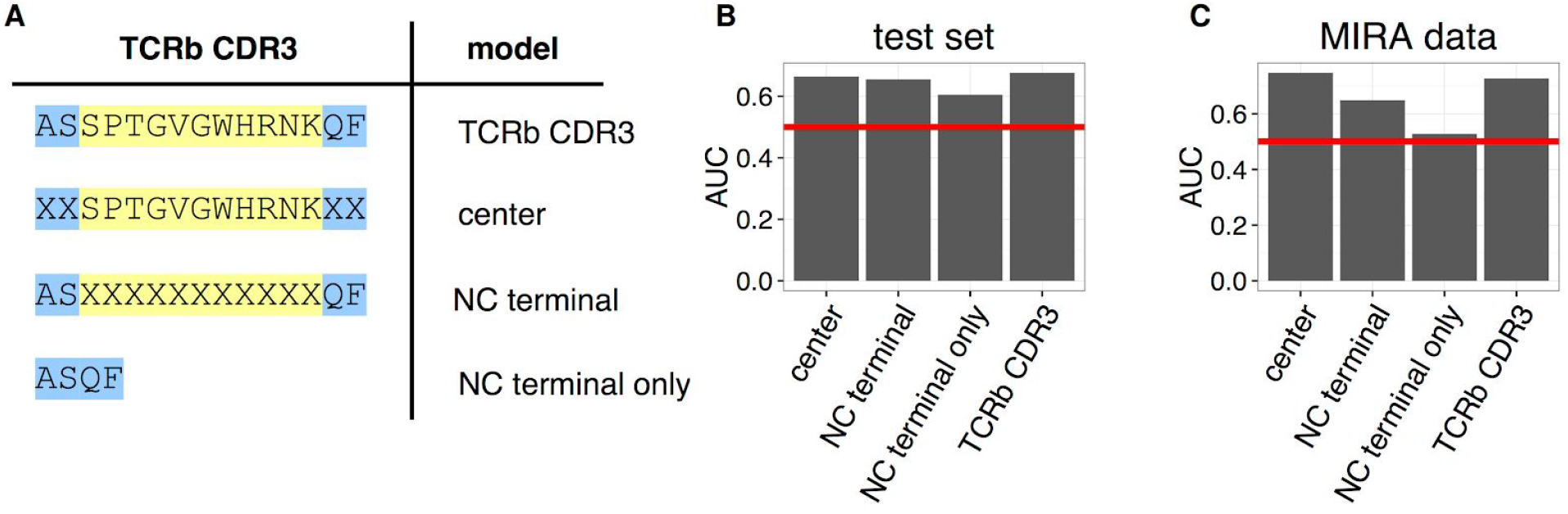
A) To investigate V+J gene bias, models with different representations of the TCRb CDR3 sequence were trained. B+C) Performance of the different models on the test and MIRA datasets given in AUC, the red line denotes random performance.

### Experimental validation of the model

Next, we set out to validate our model in a real-life biological setting. T cells from four donors were sorted into a positive subset, containing TCRs responsive to the four HLA-A*02:01 restricted peptides (GILGFVFTL, GLCTLVAML, NLVPMVATV, YVLDHLIVV) and a negative subset, containing TCRs responsive to 92 other HLA-A*02:01-restricted peptides (table S3). The CDR3 sequences were obtained from the TCR beta chains of each of these subsets. In parallel, we trained a model on the combined IEDB and MIRA data. This combined data set has a large amount of TCR data for the four positive peptides (GILGFVFTL, GLCTLVAML, NLVPMVATV and YVLDHLIVV), and we would hence expect to be able to predict the interactions to one or more of the peptides for the TCRs in the positive subset and lack of interaction for the TCRs in the negative subset. To validate that this was indeed the case, we predicted the interaction between each TCR and the four positive peptides and identified the maximum prediction value for each TCR among those four peptides. Figure 5 compares these maximum prediction values for the positive and negative samples, showing higher prediction values for the TCRs that are able to recognize one of the four peptides (p-value < 0.01, student T-test). In line with what we observed in Figure 3B, the positive TCRs achieve a median prediction value close to 0.25, while the vast majority of negative TCRs result in predictions below 0.25. 67 of the 314 unique positive TCRs are identical with TCRs in the IEDB and MIRA data sets, likely representing public TCR sequences. No identical TCRs can be found between the negative fraction and IEDB or MIRA data.

**Figure 5:**
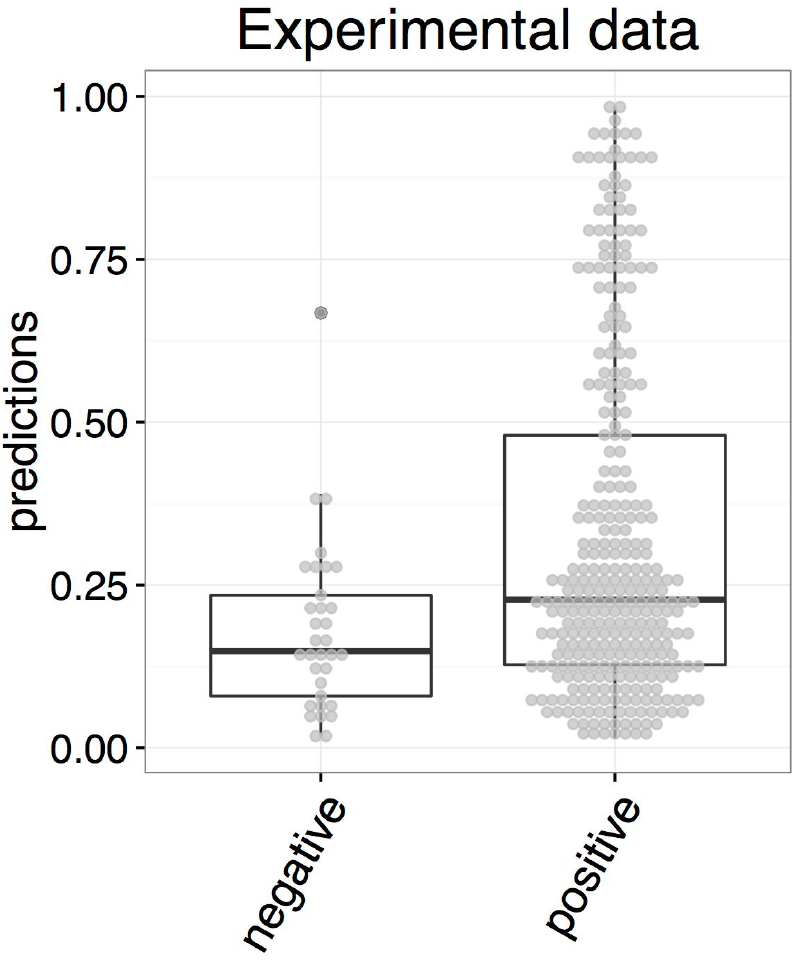
Prediction values for TCRs recognizing one of four tested peptides (positive) and TCRs not recognizing any of the tested TCRs (negative).

## Discussion

We present a model capable of predicting the cognate target of a given TCR based on the amino acid sequences of the peptide and CDR3 region of the TCR beta chain. The underlying model is a convolutional neural network (CNN). The performance of the model was evaluated in several benchmarks demonstrating a high ability both to separate T cell receptors specific for the set of peptides included in the training data from T cells specific to irrelevant peptides, and to identify the correct cognate target for a given TCR.

It has been suggested previously that learning the rules of TCR antigen recognition is extremely difficult since TCRs can rearrange themselves upon contact with the peptide-MHC complex (38) and thereby gain immense cross-reactivity (39, 40). While the necessity and role of TCR cross-reactivity in the immune systems function remains to be elucidated (41), growing evidence suggests that T cell receptors specific to a common target share common properties (17, 18). Increasing amounts of data are now available linking TCR sequences to their cognate targets. The predictive power of the model proposed here is in line with these observations.

Given the fact that most TCR sequencing projects focus on characterizing the CDR3 region of the T cell receptor beta chain sequence, we chose to train a model based on this part of the TCR only. It is clear that this potentially has limited the predictive power of our model, and that future extension of the model would benefit from being trained on paired T cell sequence data covering both the alpha and beta chain and possibly including information describing the V and J genes of the rearrangement.

The training data used here was obtained from the IEDB. Alternatively this data could have been obtained from the VDJdb, another database curating and providing TCR sequences and their cognate targets published in literature. Both databases are an extremely valuable resource to the community.

When aiming to train a model to predict TCR specificity, negative data is needed but not readily available in resources such as the IEDB and VDJdb. This is because the underlying experiments identify interacting TCRs and do not specifically report non-interacting TCRs. Our approach to resolve this, was to make mismatching combinations between peptides and TCR sequences, keeping the frequency of peptides in positive and negative data equal. The advantage of this approach is that the model is prevented from simply learning interacting peptides or TCRs by heart, due to the equal amount of positive and negative examples.

We envision that our model would likely be used to filter out TCRs from repertoire sequencing that are able to interact with a given peptide. This task of identifying very few sequences out of a pool of many requires a model of great specificity. To increase the specificity of our model it was necessary to add more TCR sequences and peptides as negative examples to the training. The negative TCR sequences were obtained by repertoire sequencing of healthy individuals and paired with the peptides in the IEDB and further self peptides identified by ligand elution assays to be presented by HLA-A*02:01. Combining TCR sequences of healthy individuals with human self peptides should result in true negative examples in the vast majority of cases, but when combining those TCRs with the peptides present in the IEDB data, which are to a large extent well studied influenza and herpes virus epitopes (42), they might result in some false negatives. However, as the chances of combining the right TCR with the right peptide in this setting are slim, in the vast majority of cases we expect to obtain true negatives with this approach. In the case of the MIRA data, the experimental setup requires TCRs to be specific to only one of the 16 assayed peptides, hence eliminating the possibility of discovering cross-reactive TCRs (15).

We chose to set up our model as a CNN. This type of model is highly flexible and has earlier been demonstrated to be highly suited to discover motifs in unaligned input sequences of varying length (28, 29). Additionally convolutional networks are often faster and easier to train than recurrent long short-term memory networks (LSTMs). Future work will tell if improved predictive power can be obtained combining different network architectures (43).

Currently available datasets of TCR-peptide interactions contain many more TCR sequences than peptides and the peptides share very limited similarity in general. It is therefore expected that a model trained on these data will have limited power to extrapolate predictions to unknown peptides. This is also what we observed when evaluating the performance of the model on peptides not present in the IEDB training data. Given these observations, we recommend that the model should presently only be used to make predictions for the four peptides covered by abundant TCR data in the combined IEDB and MIRA data.

Due to the limited amount of TCR donors in the current data set, it is a concern that models might overfit to a bias in V and J gene usage in the donors. To investigate the extent of this in our data set, we trained different models masking the N- and C-terminal regions or the center of the TCR beta CDR3 regions. We found that we could indeed train models that performed well on the test set when supplying information about the N- and C-terminal regions and the length of the CDR3 loop. When partitioning the data sets, we did not account for reducing redundancy between partitions based on the V and J genes, we only compared the entire CDR3 sequences in our approach to redundancy reduction. Therefore the same bias in V and J gene usage is likely present throughout all partitions, explaining the high performance on the test set. When evaluating models on the independent MIRA data set, obtained from a different set of donors, we find that only models trained on the central part of the CDR3 sequence generalize well. This indicates that there is indeed a bias in V and J gene usage to which a model can overfit, but there is also a signal in the central part of the CDR3 region, defining the specificity of the TCR which is consistent across several data sets and donors.

Substantial efforts have recently been dedicated to elucidate what properties of a TCR dictate its specificity, and publications suggest TCRs sharing a common target are characterized by sharing, to some degree, a common motif (17, 18). This is also the underlying assumption of the model presented here. To further validate our approach, we performed a TCR sequencing experiment where we obtained two sets of TCRs: one specific to the four peptides covered by our model, and one non-specific to any of these peptides. As expected, the model achieved higher interaction predictions for TCRs recognizing one of the four known cognate target peptides. More detailed results could have been achieved by determining the exact specificity of each TCR. This was however not possible due to limited funds.

In conclusion, we have successfully trained a model to predict interactions between TCRs and their cognate, HLA-A*02:01 restricted peptide target. Our results indicate that accurate prediction based only on the TCR beta chains CDR3 region and amino acid sequence of the peptide is feasible. Due to the small amount of training peptides, the model can however at present only be applied to the limited set of peptides included in the training data. However as more data becomes available, we expect the predictive power of the model to increase, and allow for accurate predictions also for uncharacterized peptides as has been observed earlier for the pan-specific prediction models of peptide-MHC interactions (44). Finally, the presented model framework is highly flexible and allows for the straight forward integration of the MHC molecule or TCR alpha chain in the future when data becomes available, to train a truly global prediction method.

## Aknowledgements

We thank Michael Schantz Klausen for assistance in setting up the webserver.

## Supplementary

**Figure S1:**
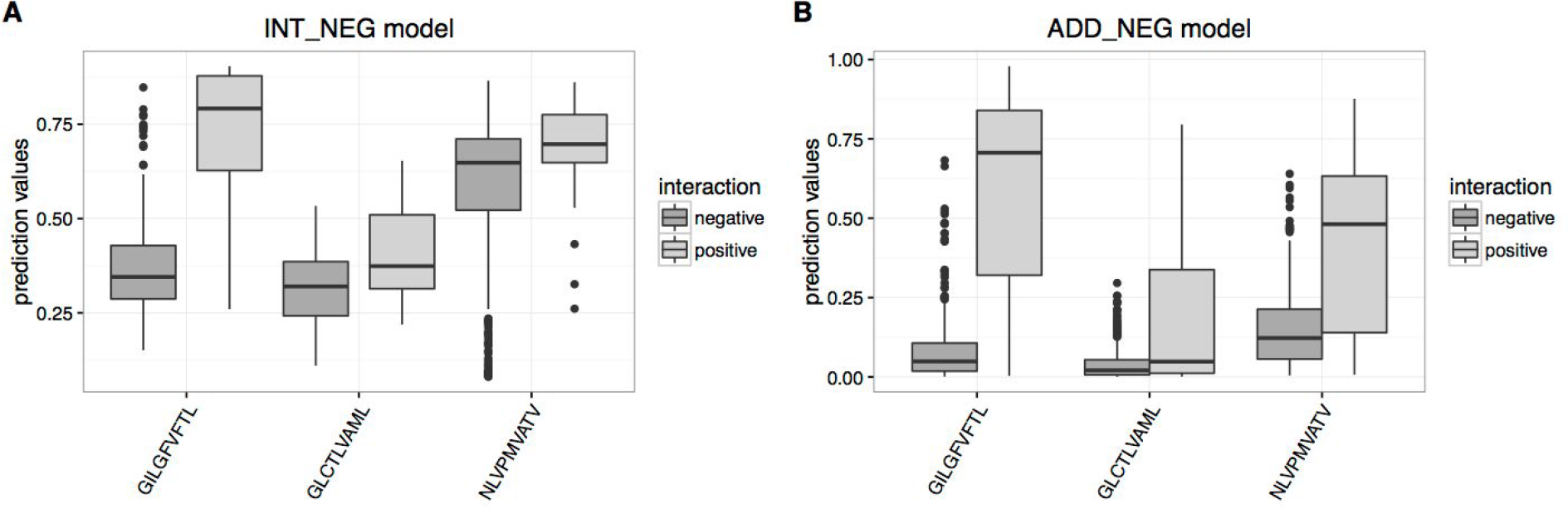
Prediction values for each TCR-peptide combination among the 3 peptides in the evaluation data and all MIRA TCRs. A) IEDB internal negative data model B) IEDB model trained with additional negative data.

**Table S1:**
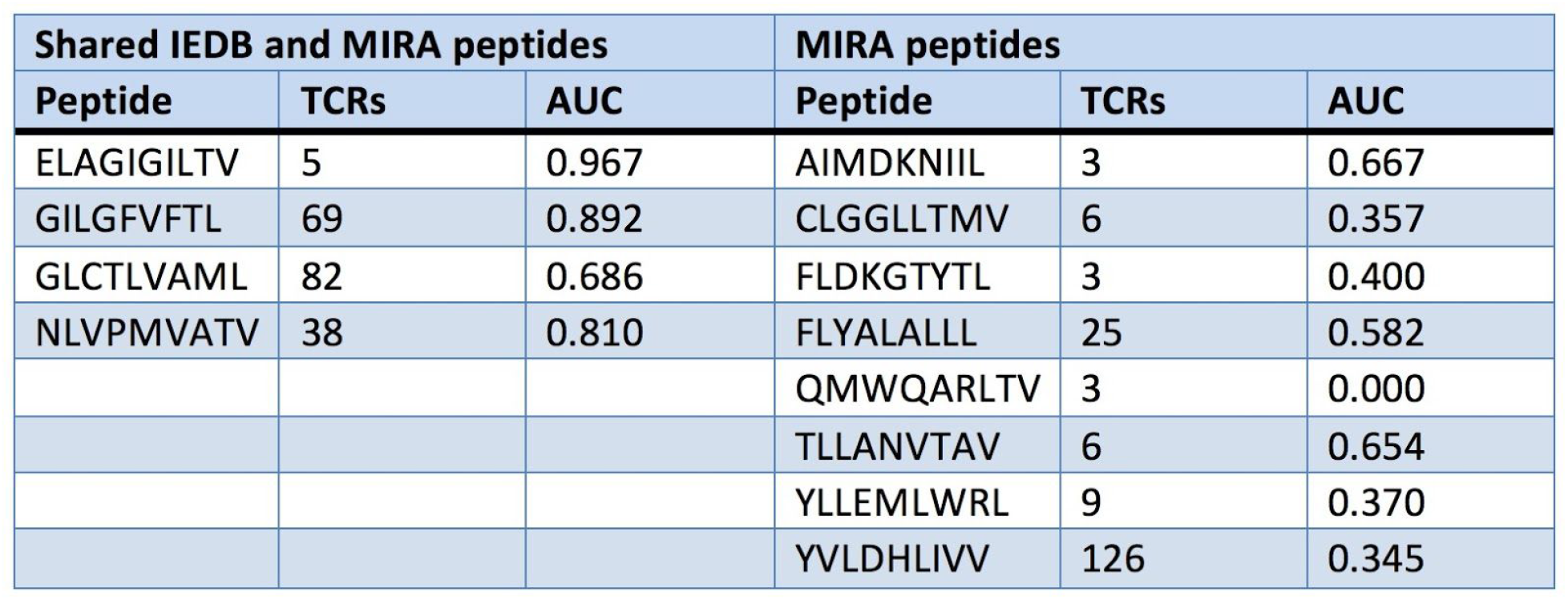
AUC per peptide of the MIRA data predicted with a model trained on IEDB with additional negative data.

**Table S2:**
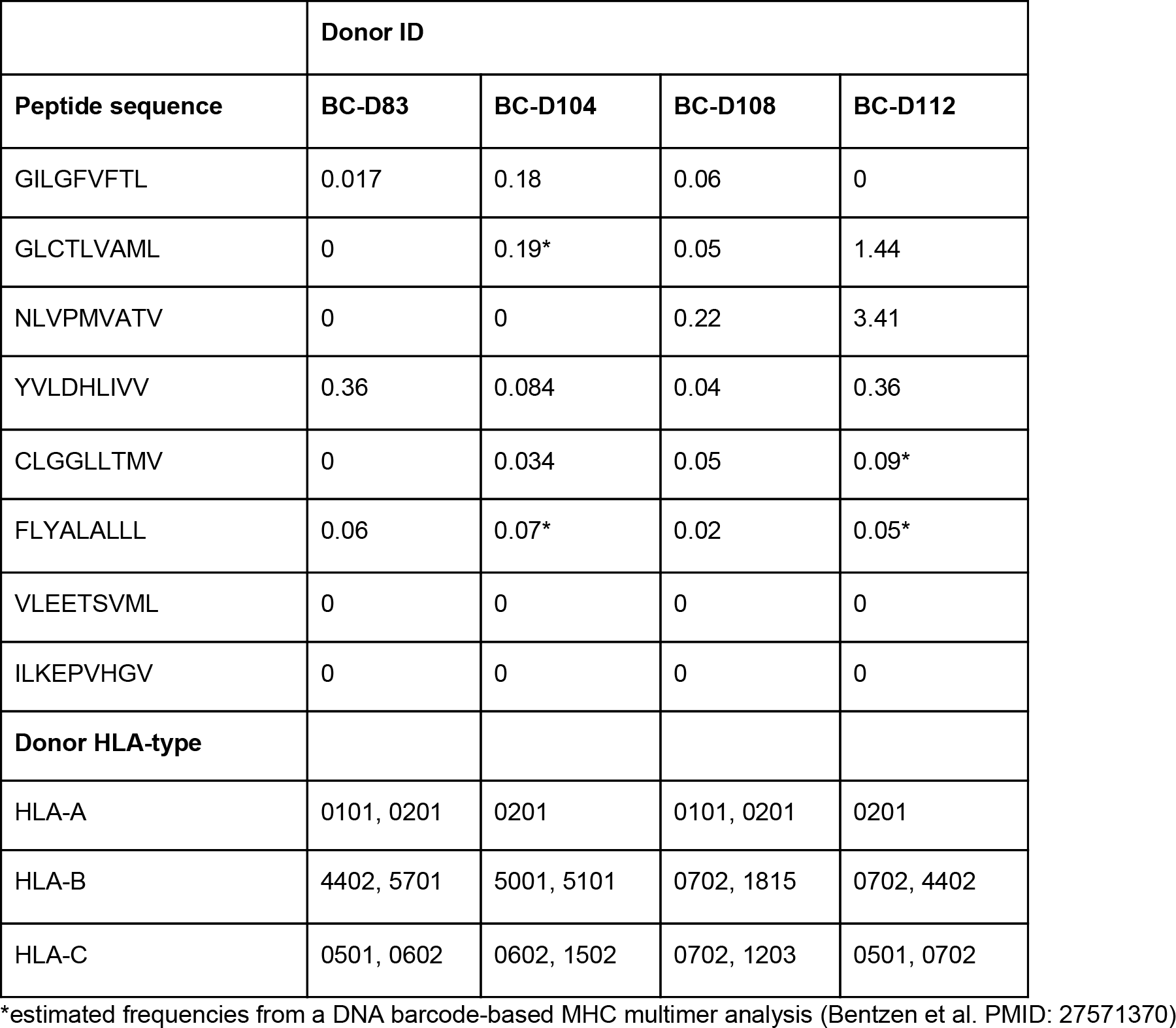
Previously detected responses (% of CD8 T cells).

**Table S3:**
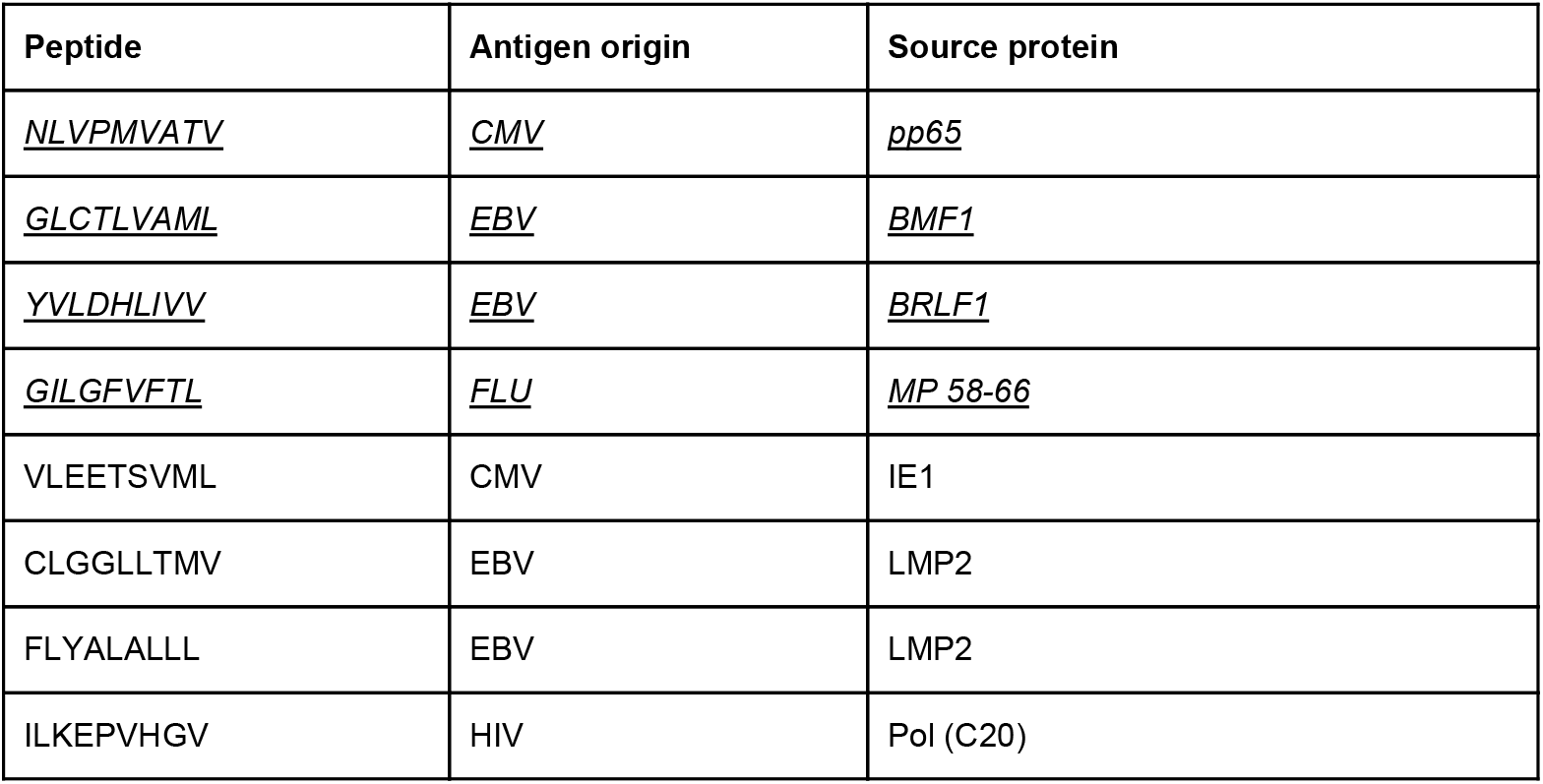
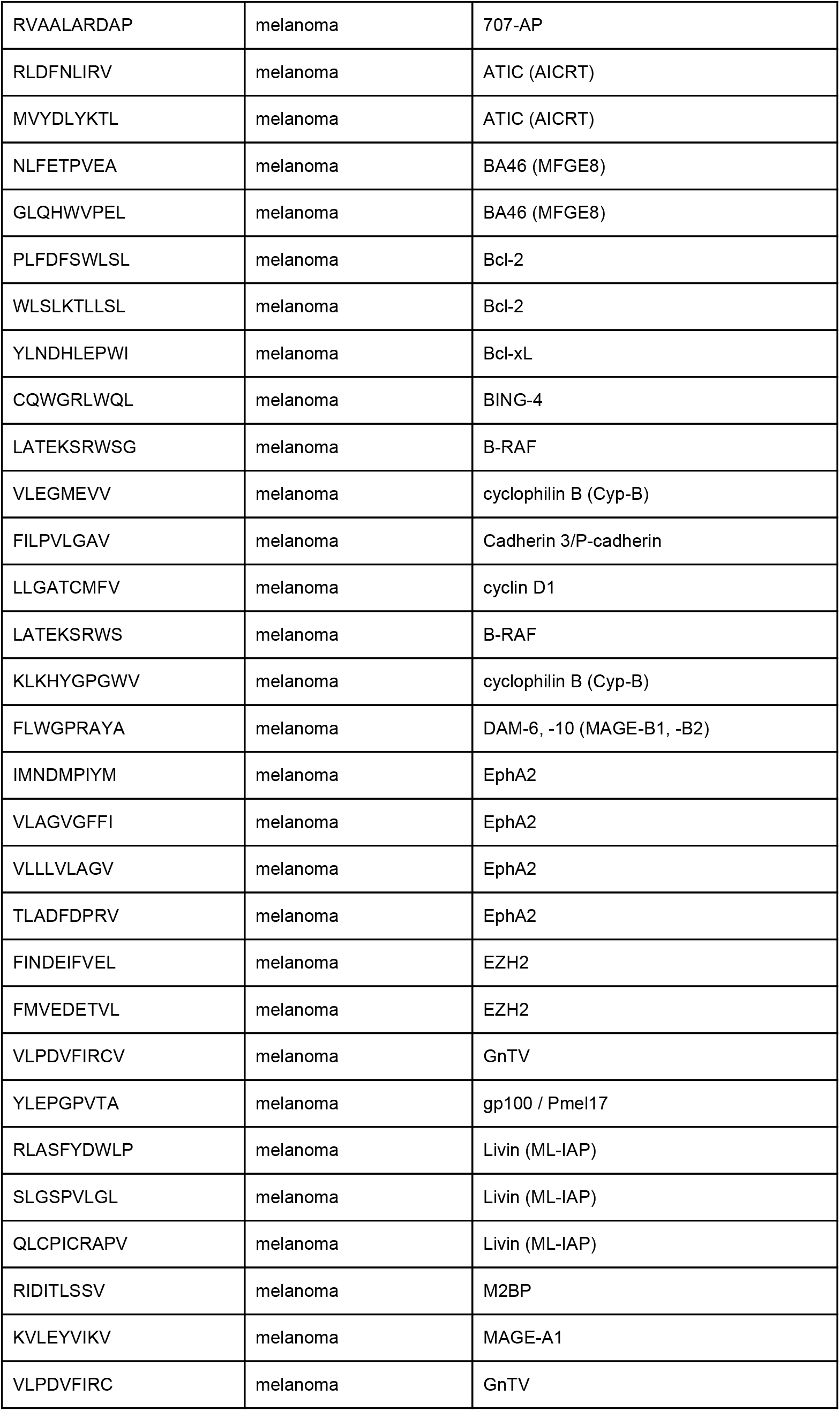
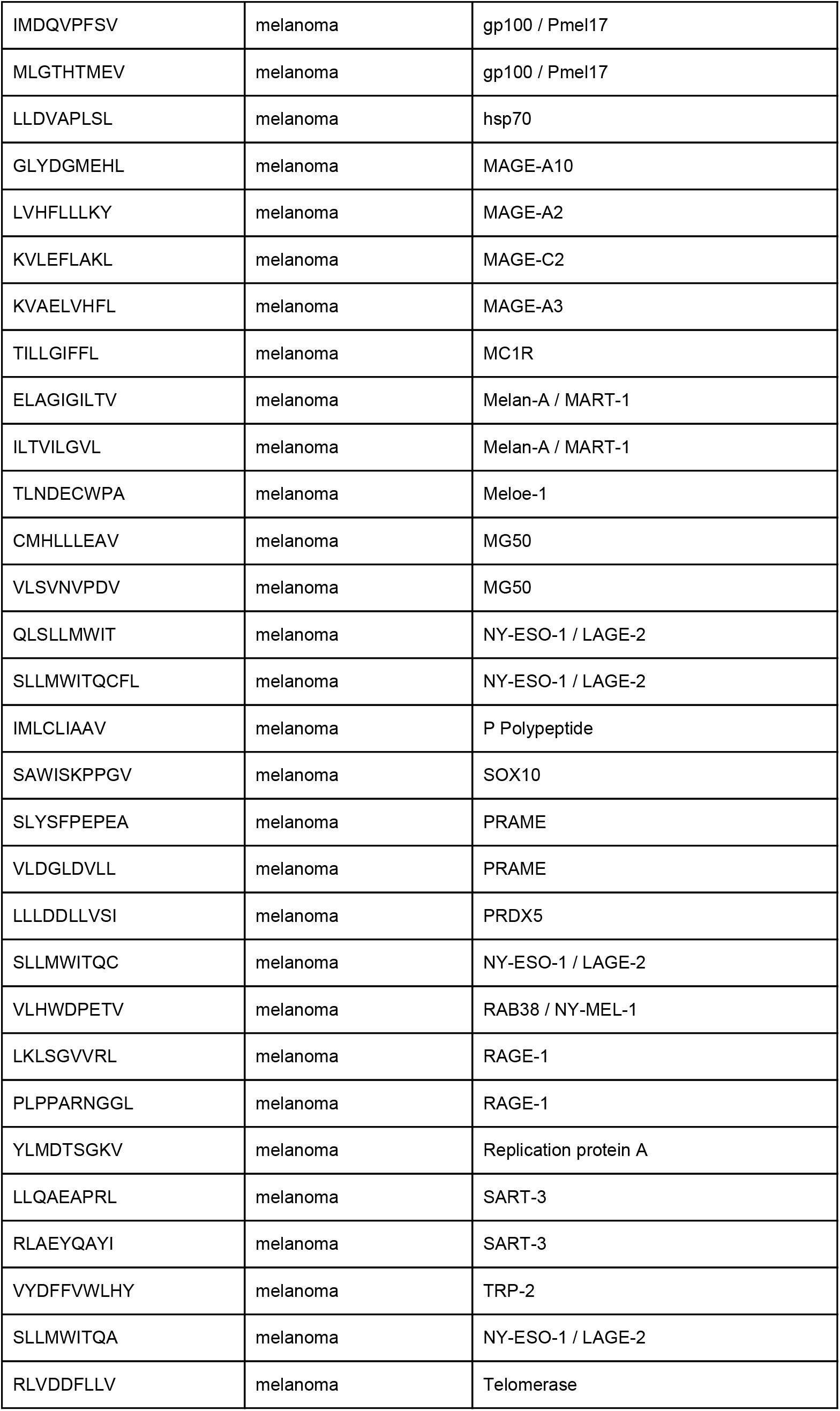
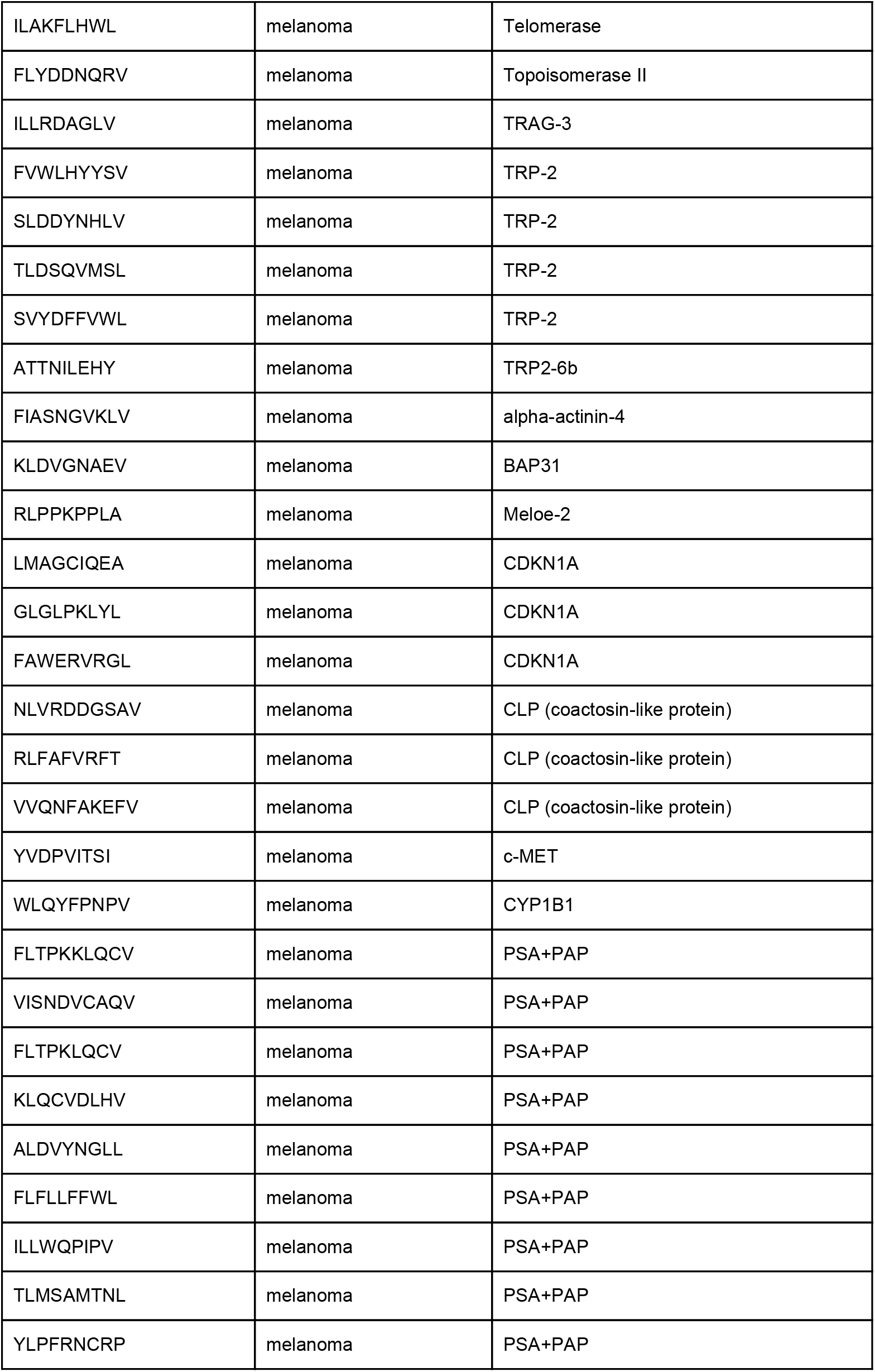
List of peptides used to isolate TCR sequences. All peptides are HLA-A*02:01 restricted.

